# Neural Tracking Measures of Speech Intelligibility: Manipulating Intelligibility while Keeping Acoustics Unchanged

**DOI:** 10.1101/2023.05.18.541269

**Authors:** I.M Dushyanthi Karunathilake, Joshua P. Kulasingham, Jonathan Z. Simon

## Abstract

Neural speech tracking has advanced our understanding of how our brains rapidly map an acoustic speech signal onto linguistic representations and ultimately meaning. It remains unclear, however, how speech intelligibility is related to the corresponding neural responses. Many studies addressing this question vary the level of intelligibility by manipulating the acoustic waveform, but this makes it difficult to cleanly disentangle effects of intelligibility from underlying acoustical confounds. Here, using magnetoencephalography (MEG) recordings, we study neural measures of speech intelligibility by manipulating intelligibility while keeping the acoustics strictly unchanged. Acoustically identical degraded speech stimuli (three-band noise vocoded, ∼20 s duration) are presented twice, but the second presentation is preceded by the original (non-degraded) version of the speech. This intermediate priming, which generates a ‘pop-out’ percept, substantially improves the intelligibility of the second degraded speech passage. We investigate how intelligibility and acoustical structure affects acoustic and linguistic neural representations using multivariate Temporal Response Functions (mTRFs). As expected, behavioral results confirm that perceived speech clarity is improved by priming. TRF analysis reveals that auditory (speech envelope and envelope onset) neural representations are not affected by priming, but only by the acoustics of the stimuli (bottom-up driven). Critically, our findings suggest that segmentation of sounds into words emerges with better speech intelligibility, and most strongly at the later (∼400 ms latency) word processing stage, in prefrontal cortex (PFC), in line with engagement of top-down mechanisms associated with priming. Taken together, our results show that word representations may provide some objective measures of speech comprehension.

**Significance Statement:** Electrophysiological studies have shown that brain tracks different speech features. How these neural tracking measures are modulated by speech intelligibility, however, remained elusive. Using noise-vocoded speech and a priming paradigm, we disentangled the neural effects of intelligibility from the underlying acoustical confounds. Neural intelligibility effects are analyzed at both acoustic and linguistic level using multivariate Temporal Response Functions. Here, we find evidence for an effect of intelligibility and engagement of top-down mechanisms, but only in responses to lexical structure of the stimuli, suggesting that lexical responses are strong candidates for objective measures of intelligibility. Auditory responses are not influenced by intelligibility but only by the underlying acoustic structure of the stimuli.

## Introduction

When we listen to speech, our brains rapidly map the acoustic sounds into linguistic representations while recruiting complex cognitive processes to derive the intended meaning (1, 2). A fundamental goal in auditory neurophysiology is to understand how the brain transforms the acoustic signal into meaningful content. Along this avenue, a large body of research has demonstrated that neural responses time lock to different features of the speech signal (“neural speech tracking”) (2, 3). These features have primarily included acoustic features like the speech envelope and envelope onset, but more recently also include linguistic units such as word onsets, phoneme onsets and context-based measures along different levels of the linguistic hierarchy. However, it remains unclear how these neural tracking measures are affected by intelligibility.

Neural tracking measures of intelligibility have often been investigated using experimental designs that manipulate intelligibility by altering the underlying acoustical structure, such as through time compression (4, 5), disruption of spectro-temporal details (6–8) and time reversal (4, 9). While these studies successfully demonstrate differences in cortical tracking responses, interpreting these findings is not straightforward, as the observed changes in the neural response may arise from the alterations in the acoustic waveform (bottom-up), rather than the intelligibility change itself (top-down). Intelligibility related neuro-markers derived from neural responses play a crucial role in advancing our understanding of the neurophysiology of the speech understanding. They would contribute to the clinical evaluation of auditory function across diverse clinical populations and aid in the hearing device evaluation (10). In situations where obtaining verbal responses are challenging, such as with infants or individuals with cognitive disabilities, as well as when subjective estimates are affected by individual differences, neuro-markers of intelligibility would offer a non-invasive and objective means to investigate the underlying neural processes.

In this study, we disentangle intelligibility from any underlying acoustical confounds by using single instances of noise-vocoded speech combined with a priming paradigm. Noise-vocoded speech greatly reduces intelligibility by removing spectral details but still preserving the slow temporal envelope, and is often used as a surrogate for speech perception by cochlear implant patients (11). It is generated by processing the original speech signal through a multi-frequency-channel vocoder, where higher numbers of channels retain more intelligibility due to the retention of more spectral information. With a sufficiently low number of channels, most vocoded speech is largely unintelligible without practice. In the current study, we used three-band noise-vocoded speech. Magnetoencephalography (MEG) data were recorded from young adult participants (*N* = 25) as they listened to a passage of noise-vocoded speech, first before any priming (PRE), followed by listening to the original, non-degraded version of the same passage to invoke priming (CLEAN), and then finally listening to the same noise-vocoded speech passage as before (POST), repeated for 36 trials (see Figure 1(A)). At the end of each vocoded speech passage, participants were asked to rate the perceived speech clarity on a scale from 0 to 5. Compared to previous studies (12–15) using a similar paradigm with shorter sentences (<5 s), the current study uses much longer passages (∼20 s), making it challenging to rely solely on short-term memory to understand POST vocoded speech.

**Figure 1.**
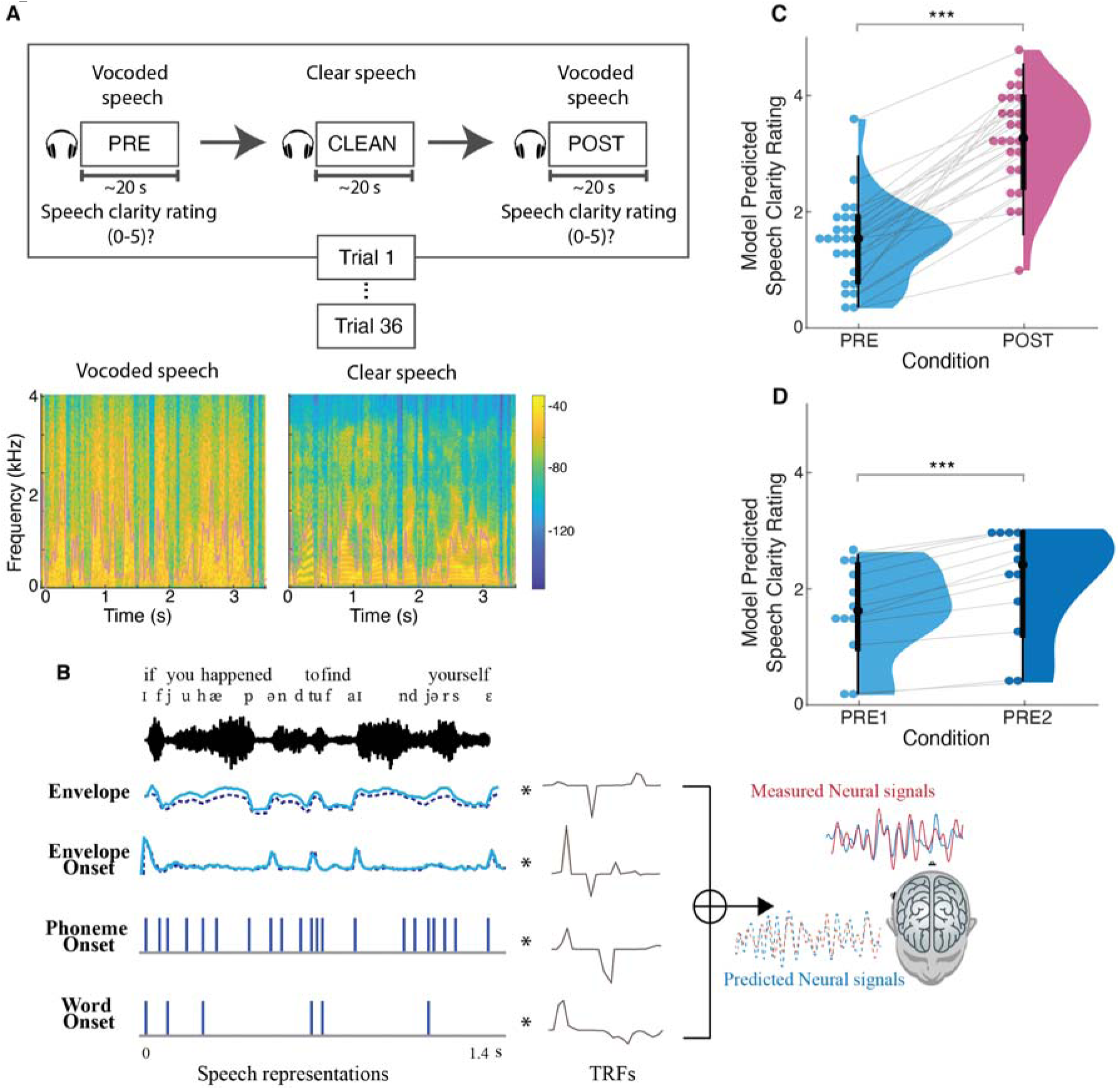
Experimental methodology, spectro-temporal characteristics of the speech stimuli, analysis framework, and behavioral results. **(A)** (top) Participants listened to 36 audiobook passages in vocoded and clear speech conditions. Each trial consisted of presentation of the same passage in vocoded format (PRE), clear format (CLEAN) and identical vocoded format (POST). At the end of each vocoded speech passage, participants rated the speech clarity on a scale from 0 to 5. (bottom) Spectrograms and temporal envelopes (overlaid in pink) of vocoded and clear speech. Most spectral and fine temporal details of the vocoded speech are lost (e.g., the pronounced vertical striping), but the broad temporal envelopes of the clear and vocoded speech are very similar. **(B)** Multivariate temporal response function (mTRF) analysis of MEG and predictor variables illustrated with sample of stimuli. Individual TRFs represent the brain’s responses to the corresponding speech representations at different time lags. (C) Linear Mixed Effects Model (LMEM) predicted speech clarity ratings (0–5) for PRE and POST vocoded conditions. Perceived clarity of the vocoded speech is significantly improved after the clear speech priming. **(D)** LMEM model predicted speech clarity ratings for PRE1 and PRE2 vocoded conditions in the control study. In the control study, subjects listened to the vocoded speech passages without priming, where the two vocoded speech presentations are denoted by PRE1 and PRE2. Perceived clarity of the second presentation is enhanced compared to first presentation, but the improvement is smaller compared to that with priming. *p<0.05, **p<0.01, ***p<0.001

To characterize how different speech features are tracked in the cortical response, we utilized multivariate temporal response function (mTRF) analysis (16–18), as illustrated in Figure 1(B). Similar to conventional event-related potentials (ERPs), TRFs are used to assess how the brain reacts to different speech features over time. In contrast to ERPs, which rely on averaging numerous short responses to determine the brain’s reaction to given stimulus, TRF enables the examination of brain’s responses to continuous speech, as well as simultaneous responses to multiple speech features (1). We included three classes of speech features for which to extract responses: acoustic, sub-lexical, and lexical. The specific features employed were speech envelope, envelope onset, phoneme onset, and word onset, to cover a range of neural responses from acoustic processing to lexical level processing.

We first determined whether each of these features are represented in the cortical response by evaluating the explained response variability for each of the speech conditions (PRE, POST and CLEAN). Then we investigated how the cortical representations of the different speech features are modulated by intelligibility (PRE vs POST) and by acoustics (vocoded PRE and POST vs CLEAN) at different auditory processing stages, by comparing the TRF peak amplitudes and explained variability. To further test whether differences in PRE vs POST may be attributable to intelligibility, as opposed to mere passage repetition, a control study was conducted involving 12 subjects who listened to the same passages but in a different order without priming (i.e., PRE1, PRE2, CLEAN). All analysis were performed on source-localized brain responses, and were restricted to temporal, frontal and parietal brain regions.

## Results and Discussion

### Behavioral performance increases with speech priming

We first determined the extent to which the speech priming (perceptual learning) affected the speech intelligibility between PRE and POST vocoded conditions using perceived speech clarity ratings. A linear mixed effect model (LMEM) was modelled with perceived clarity rating as the dependent variable, condition as a fixed effect, and random intercept and slopes for condition by subject as random effects (*clarity rating ∼ 1 + condition + (1+condition|Subject)*). As illustrated in Figure 1(C), the fixed effects of condition revealed that rated speech clarity in the POST vocoded condition is improved compared to PRE (increase = 1.81, *SE* = 0.16, *p* < 0.001). In the control study (Figure 1(D)) where the CLEAN speech was only presented after the second vocoded speech presentation, perceived clarity ratings did show significant improvement in clarity even without the clean speech priming (increase = 0.52, *SE* = 0.10, *p* < 0.001). However, the effect was substantially smaller than the priming effect (PRE2-PRE1 vs POST-PRE = -1.28, *p_perm_* < 0.001), suggesting that priming has a substantially larger impact on speech clarity relative to acoustical learning without priming.

This result supports the idea that presentation of the clear speech, which provides information regarding both the linguistic content (i.e., words, content) and physical acoustical structure (i.e., rhythm, pace) of the degraded speech, facilitates (a top-down influence on the) understanding of POST vocoded speech. Thus, in agreement with previous studies, speech that is acoustically identical but initially unintelligible can be made intelligible through perceptual learning (12, 14, 15, 19–23)

### Neural responses to acoustic features do not index speech intelligibility, only acoustics

To test the extent to which each of the acoustic features are represented in the brain and for each condition, we first compared the predictive power, measured as the explained variability (*R^2^*) of the full model against a reduced model that excluded the predictor of interest (see Figure 2(B,C) right column, brain plots). This prediction accuracy analysis revealed that both acoustic envelope and envelope onset significantly contribute to the model’s absolute predictive power for all three conditions, PRE (envelope: *t_max_* = 6.5, *p* < 0.001, onset: *t_max_* = 6.2, *p* < 0.001), POST (envelope: *t_max_* = 7.3, *p* < 0.001, onset: *t_max_*= 5.9, *p* < 0.001) and CLEAN (envelope: *t_max_* = 6.3, *p* < 0.001, onset: *t_max_* = 7.24, *p* < 0.001). These findings suggest that acoustic features are processed irrespective of the stimuli intelligibility. The anatomical distribution of significant acoustic feature processing for each condition was observed in locations spreading spatially from Heschl’s gyrus (HG) to superior temporal gyrus (STG) and much of temporal lobe. This distribution was bilateral, and also dominantly in the right hemisphere except for clean speech envelope and vocoded speech envelope onset processing ((left vs right hemisphere) envelope: PRE_*t_max_* = -5.35, *p* = 0.02, POST_*t_max_* = -5.25, *p* = 0.004, CLEAN_*t_max_* = -3.3, *p* = 0.41, envelope onset: PRE_*t_max_* = - 4.27, *p* = 0.02, POST_*t_max_*= -3.46, *p* = 0.39, CLEAN_*t_max_* = -4.68, *p* = 0.006). This pattern of source localization, including right-hemisphere dominance, suggests that the processing of these speech features relies heavily on bottom-up driven mechanisms (24, 25).

**Figure 2.**
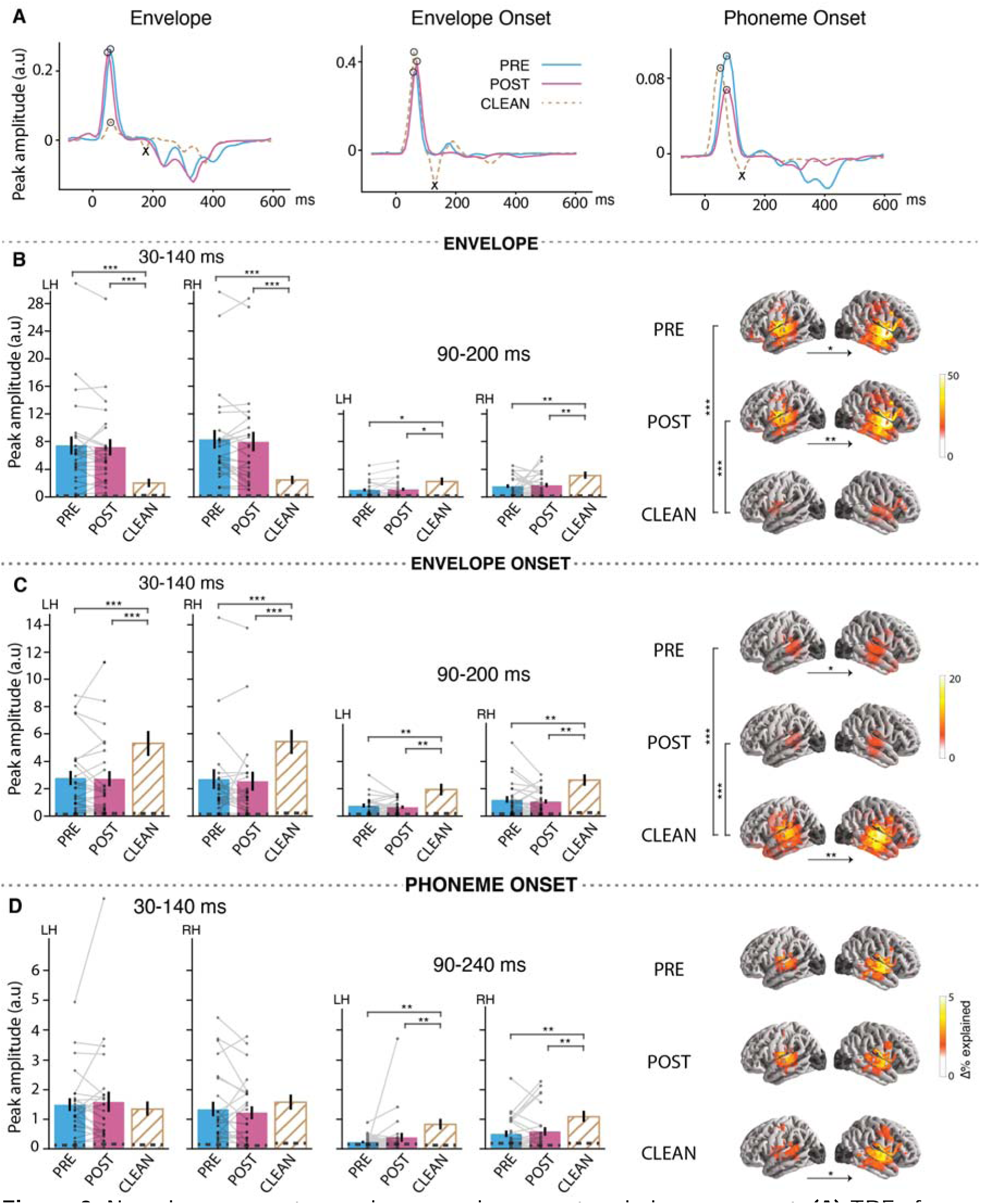
Neural responses to envelope, envelope onset and phoneme onset. **(A)** TRFs from a representative subject in source space, visualized as a single time series using principal component analysis (PCA). The TRFs exhibit an early peak (positive polarity peak marked by) and late peak (negative polarity peak marked by). **(B, C, D)** (left) Bar plots (mean±standard error (SE)) compare the measured peak amplitudes in the denoted time window (for left (LH) and right (RH) hemispheres separately) for envelope, envelope onset and phoneme onset respectively. Individual subject data points are shown for both vocoded conditions with each subject’s data points connected by lines (see supplemental Figure S1(A) for CLEAN speech individual data points). Peak amplitudes were extracted as the maximum peak of the sum of absolute current dipole strengths across sources with a specific polarity, where the polarity was determined from the current directions from the original source TRFs. (right) brain plots show cortical regions where the given speech feature significantly improves the model fit over and beyond other speech features in the model. Vertical significance brackets indicate significant differences between conditions, and horizontal significance brackets below brain plots indicate lateralization differences. The dashed lines within the bars represent the noise floor, where peaks for a noise model were extracted using the same steps as above. Significant differences were found between vocoded vs CLEAN passages, but no differences were observed between PRE vs POST.

In order to determine whether the processing of acoustic features differed based on speech intelligibility (PRE vs POST) or acoustics of the stimuli (vocoded vs clean), we compared the model improvements between speech conditions for each acoustic feature individually. Pairwise mass-univariate related samples *t*-test revealed that there is no significant difference between PRE vs POST vocoded speech with respect to both envelope (*t_max_*= -3.9, *p* = 0.57) and envelope onset (*t_max_* = 4.39, *p* = 0.09). In contrast, the variance explained due to envelope processing was significantly stronger for vocoded speech compared to clean speech (*t_max_* = 6.8, *p* < 0.001), and the opposite for envelope onset processing (*t_max_*= -7.44, *p* < 0.001), suggesting that auditory responses are mainly driven by the manipulations in the stimuli.

We then investigated how the brain responds to acoustic speech representation at different cortical processing stages with an amplitude analysis of the TRF waveforms. In analogy to the ERP P1-N1 peaks at the corresponding latencies, the envelope TRF showed two main peaks (Figure 2(A)), a positive polarity peak at ∼50 ms latency followed by a negative polarity peak ∼100 ms, known as the M50_TRF_ and M100_TRF_ respectively (1, 17). Analogous to these envelope peaks, envelope onset responses also showed two main peaks at ∼75 ms and ∼130 ms, consistent with previous studies (1, 26). The M50_TRF_ and M100_TRF_ peaks can be ascribed to different auditory cortical processing stages with the corresponding latencies (27). It has been suggested that the early M50_TRF_ peak dominantly reflects the neural encoding of low-level processing, e.g., physical acoustic of the stimuli (28, 29), whereas the M100_TRF_ peak reflects additional higher-level processing, e.g., selective attention (17, 28, 30). Using a paired-samples *t*-test, we investigated the extent to which these peak amplitudes are affected by the intelligibility and acoustic characteristics of the stimuli (Figure 2(B,C)). In line with the prediction accuracy results above, we failed to find an effect of intelligibility for the neural responses to acoustic features (both envelope and envelope onset) for both the M50_TRF_ (envelope: *p* = 0.63, envelope onset: *p* = 0.80) and M100_TRF_ peaks (envelope: *p* = 0.71, envelope onset: *p* = 0.29)), but TRF peak amplitudes were significantly affected by the differences in acoustics for both the M50_TRF_ (PRE vs CLEAN: envelope *p* < 0.001, envelope onset *p* < 0.001) and M100_TRF_ peaks (PRE vs CLEAN: envelope *p* = 0.03, envelope onset *p* = 0.003); effect sizes are reported in Supplemental Table S1. These results support the previous finding that such low-frequency auditory cortical responses are not necessarily driven by the intelligibility or the linguistic content of the stimuli, but rather reflects the sensitivity to differences in sensory input (12, 15, 20, 23). Some prior studies using similar experimental paradigms have instead reported stronger envelope tracking for trials associated with better speech intelligibility. These studies used different neural tracking indices, either more complex (23, 31), more robust against low neural SNR (speech envelope reconstruction rather than TRF analysis) at the expense of less temporal resolution (13), ECoG-based high gamma responses (20) or did not yield a clear relationship (22). Differences from the current findings may be due to several factors, including differences in the neural measures employed or high gamma responses reflecting different neural sources compared to MEG/EEG. A simpler explanation, however, may be to ascribe such differences to the employment here of higher-level linguistic feature encoding (e.g., lexical segmentation responses), which allow a finer grained analysis of which aspects of the speech tracking responses are due to acoustic vs. higher-level features. As can be seen from the present results, envelope and envelope onset are very sensitive to changes in the sensory input, and any changes there associated with speech intelligibility may be very subtle. Indeed, improved data processing techniques and further refinements in acoustic neural indices might well alter uncover changes with the intelligibility for the lower-level acoustic responses. Additionally, several studies reporting increased intelligibility associated with increased cortical speech tracking relied on changing the corresponding underlying acoustics, creating a confound (6, 13).

The M50_TRF_ peak of the envelope response was significantly stronger for vocoded speech compared to clean speech, whereas this effect was reversed for the envelope onset response (Table S1), suggesting distinct mechanisms are involved in envelope and envelope onset processing. Analysis of the ratio of M50_TRF_ between envelope and envelope onset comparison across conditions provided additional support for the reliability of this reversing effect (PRE vs POST: *t_24_* = 0.53, *p* = 0.56, PRE vs CLEAN: *t_24_* = -6.25, *p* < 0.001). This finding for the early envelope TRF peak is consistent with previous studies (8, 22, 32, 33), where it was proposed that this effect is modulated by task demand (22) or sensory gain of acoustic properties (8); this early peak is too early to be modulated by attention (17). Instead, we propose that the higher envelope TRF amplitudes observed for vocoded speech are a result of its low spectral variability, which leads to higher levels of synchronization along the tonotopic axes, resulting in stronger MEG responses. In contrast, reduced vocoded speech envelope onset tracking can be attributed to the loss of salient acoustic onsets in vocoded speech, a result of the loss of spectrotemporal details intrinsic to the process of vocoding (see Methods, predictor variables).

Compared to vocoded speech, both envelope and envelope onset in clean speech exhibited stronger M100_TRF_ peak amplitudes. This suggests that the observed differences in vocoded vs clean for the M50_TRF_ and M100_TRF_ are due to different underlying brain processes, where the M50_TRF_ depends more strongly on the physical acoustics while M100_TRF_ also reflects higher-level processing (17, 28, 30). However, to the extent M100_TRF_ peak is indicative of higher-level processing, it is a striking finding that it is not influenced by intelligibility.

### Responses to Phoneme onsets

Although phoneme onsets (see Figure 2(D)) significantly contributed to the model’s predictive power in each speech condition (PRE: *t_max_* = 4.9, *p* < 0.001, POST: *t_max_* = 5.4, *p* < 0.001, CLEAN: *t_max_* = 6.8, *p* < 0.001), no significant differences were detected between the conditions. Consistent with previous studies (1), the phoneme onset TRF showed two main peaks, an early positive polarity peak ∼ 80 ms and a late negative polarity peak ∼150 ms, which were comparable to the envelope onset peaks. The amplitude of the early peak showed no difference between PRE vs POST (*p* = 0.68) nor vocoded vs clean speech (*p* = 0.28), indicating that these early responses are not modulated by either acoustics or intelligibility. However, the late negative polarity peak was stronger in clean speech compared to vocoded speech (CLEAN vs PRE: *p* = 0.003, CLEAN vs POST: *p* < 0.001, PRE vs POST: *p* = 0.31), indicating that this late processing stage is affected by the physical acoustics. Phoneme onset processing originated bilaterally from areas including primary auditory cortex and was right lateralized for clean speech (*t_max_*= -4.39, *p* < 0.02), supporting previous results that phoneme onset processing occurs early in the auditory processing hierarchy and may reflect more acoustic processing than linguistic. These results are aligned with a previous EEG study (23) that investigated the impact of perceived intelligibility on phoneme level processing (a more complex measure does show effects of priming in the delta band). Thus, despite the sub-lexical or linguistic nature of the phoneme onset feature, the neural responses suggest that, as a neural measure, it functions more like an auditory (or intermediate auditory-linguistic) measure.

### Neural responses to lexical segmentation indexes speech intelligibility

Novel in the current study, we incorporated a lexical feature to extend the investigation of the effects of intelligibility with respect to neural speech representation: the word onset response. Prediction accuracy analysis revealed that the word onset responses significantly explain additional variability in the measured neural response over and beyond acoustic and phoneme onset features, across all three speech conditions (PRE: *t_max_* = 4.6, *p* < 0.001, POST: *t_max_* = 4.6, *p* < 0.001, clean: *t_max_* = 5.1, *p* < 0.001). The word onset TRFs (see Figure 3(A)) showed two main peaks, an early positive polarity peak (∼100 ms) and a substantially later negative polarity peak (∼400 ms). The late peak is comparable to the latency and polarity of classical N400 responses (9, 34, 35), a potential marker of complex language processing, and so will be referred to here as the N400_TRF_. Interestingly, the peak amplitude comparison (Figure 3(B)) revealed that the intelligibility of the speech modulates both early (*p* = 0.02; Cohen’s d = 0.39) and late peak amplitudes (*p* < 0.001; d = 0.82), with a substantially greater effect size observed for the late peak. Comparing clean vs vocoded speech, we found that the clean speech TRF amplitudes are stronger compared to vocoded speech for both early (*p* < 0.001) and late peak responses (PRE vs CLEAN: *p* < 0.001; d = 1.20, POST vs CLEAN: *p* = 0.04; d = 0.40). Additionally, the late peak amplitude of POST is significantly closer to that of CLEAN than the corresponding amplitude of PRE is to CLEAN, suggesting that the late word onset responses of POST are more similar to CLEAN than those of PRE.

**Figure 3.**
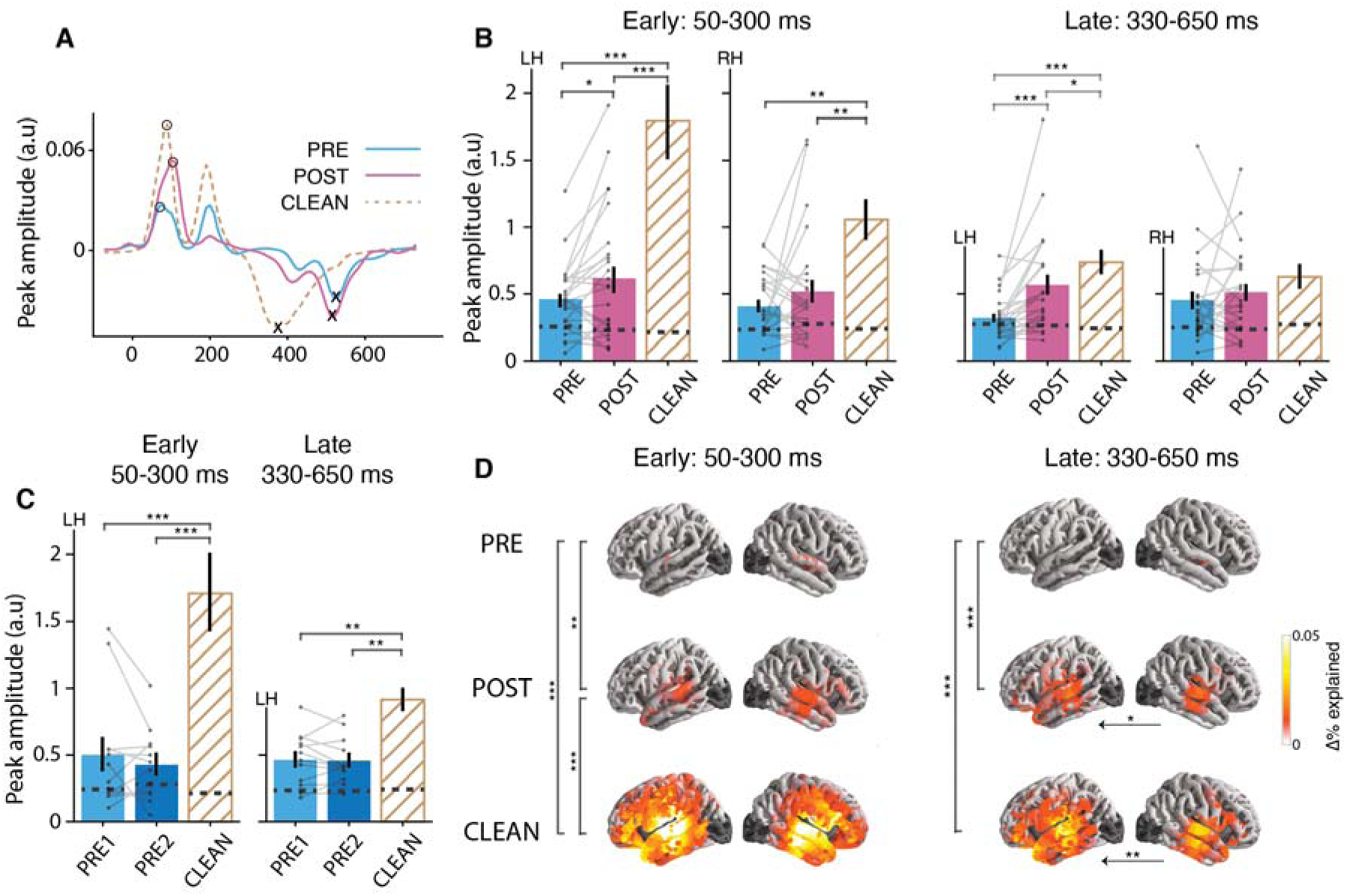
Neural responses to word onset. **(A)** The word onset TRFs for a representative subject shows two main peaks: an early positive peak (50-300 ms) and a late negative peak (330-650 ms). **(B)** Peak amplitude comparison by speech condition for early and late peaks and by hemisphere. C) Peak amplitude comparison by speech condition for early and late peaks in the control study. **(D)** Word onset contributions to the prediction accuracy, but separated for the early and late processing stages. Other details as in Figure 2. TRF peak amplitude comparison and prediction accuracy comparisons show that both early and late word onset responses are modulated by intelligibility. At the early processing stage neural responses in POST compared to PRE are stronger in STG and towards late processing stage this effect was extended to left PFC. Peak amplitude enhancement is not observed for mere passage repetition (in the control study) in either early or late responses.

Because prediction accuracy, as implemented above, integrates over a longer time window (–200 - 800 ms), it is not able to disentangle specific contributions to the prediction accuracy and source localizations of each processing stage. To address this, we conducted a separate analysis on the explained variability, focusing on early (50-300 ms) and late (330-650 ms) processing stages separately (see Figure 3(B,D)). As can be seen from Figure 3(D), prediction accuracy comparisons between PRE vs POST revealed that, during the early processing stage, neural processing is significantly stronger in POST compared to PRE in the STG (*p* = 0.01). Notably, at the late processing stage this effect was extended to the left prefrontal cortex (PFC) (*p* < 0.001). Additionally, the comparison between vocoded and clean speech revealed that, at the early processing stage, clean speech elicited stronger neural responses across much of the temporal lobe (*p* < 0.001) and towards the late stage, significant differences were confined primarily to both left STG and PFC (*p* < 0.001).

This key finding suggests that responses to word onset can serve as an index of speech intelligibility independent of the acoustics. Neural tracking of word onsets represent both bottom-up and top-down processes (18, 36, 37). The early word onset peak may dominantly reflect bottom-up driven mechanisms such as acoustics at word boundaries and automatic word segmentation, but nevertheless does show significant changes after priming. The ∼400 ms latency peak, however, is too late to be solely modulated by acoustics and could incorporate higher order word segmentation, semantic integration and other top-down processes (5, 37). When the vocoded speech is unintelligible, neither the words nor word boundaries are clear, resulting in weaker synchronized neural responses at the word onsets. As a result of priming, however, the brain has been provided with additional information (perhaps in the form of new priors) regarding the cues for words, enabling higher intelligibility and the concomitant word boundaries, enabling the emergence of word onset responses. These responses are smaller compared to those of clean speech, as might be expected due to less precise time-locking associated with still-present word boundary uncertainty. The anatomical distribution of the early processing stage, bilateral STG, may reflect a mix of both unaccounted-for auditory responses and also the expected higher-level lexical processing, while the late processing stage’s activation of left lateralized PFC may link to the engagement of top-down mechanisms. Such prefrontal activation is consistent with functional magnetic resonance imaging (fMRI) studies using a similar paradigm that found more activation in prefrontal and cingulate cortices with increased speech intelligibility (21). The current study expands on this result by leveraging MEG’s superior temporal resolution to show that these processes specifically occur in a time-locked manner: corresponding to the N400_TRF_ peak, with a ∼400 ms post-word-onset latency, and the same polarity as the N400.

### Neural responses to contextual word surprisal

One additional analysis was conducted, leveraging context-based speech representations (including those from large language models) that have recently gained popularity in the field of neural speech and language processing and may represent aspects of semantic integration and speech comprehension (2, 37–41). Contextual word surprisal was additionally included as a separate predictor, estimated using a generative pre-trained large language model (GPT-2) (42), which quantifies how surprising a word is given the previous context. Analysis revealed that contextual word surprisal responses show similar effects of intelligibility to those observed in word onset responses for the late processing stage (see Supplemental Figure S1(B)) (*p* = 0.03). The early processing stage, however, was not significantly stronger in POST compared to PRE (*p* = 0.93). The significant response to a context-based speech representation is evidence for comprehension-linked processing, in addition to and beyond mere lexical segmentation, especially at the late stage.

### Neural responses to passage repetition

Finally, to strengthen support for the idea that the observed differences in PRE vs POST word onset responses are indeed linked to intelligibility itself and are not just a side effect of passage repetition increasing familiarity, we repeated the same analysis but for the control study (Figure 3(C)). Critically, there was no word onset response change from PRE1 to PRE2 for either the early (*p* = 0.34) or late peaks (*p* = 0.74). Word onset TRF peak amplitudes were significantly larger for CLEAN speech compared to PRE1 or PRE2 (early: *p* < 0.001, late: *p* < 0.001). Furthermore, the increase in word onset TRF peak amplitudes from PRE to POST in the main study were significantly larger than changes from PRE1 to PRE2 in the control study (PRE2-PRE1 vs POST-PRE: early = -0.25, *p_perm_* = 0.02, late = -0.25, *p_perm_* = 0.006). These results add further support to the idea that, improvements in intelligibility also generate increased neural responses to lexical segmentation feature over and beyond any acoustical learning.

The null results observed in the comparison between PRE and POST for auditory and phoneme responses might potentially arise from a cancellation of enhancement and suppression effects linked with priming, prediction, and repetition (14). The control study, aimed at assessing effects of repetition without involving priming, however, did not reveal significant differences between PRE1 and PRE2 responses for both early (envelope: *p* = 0.001, onset: *p* = 0.59, phoneme onset: *p* = 0.11) and late components (envelope: *p* = 0.94, onset: *p* = 0.10, phoneme onset: *p* = 0.52), except for the early envelope response (Figure 4(A,B,C)). Early envelope responses were significantly enhanced for PRE2 compared to PRE1 (*p* = 0.001). The observed differences between envelope and envelope onset responses suggest that distinct neural mechanisms underlie these responses. Thus, while neural responses often exhibit suppression with repetition (43), this was not seen in our control study (repetition effects are typically investigated using shorter and more predictable stimuli (44), which contrast with the unintelligible and longer (∼20 s) passages used in the current study). Therefore, differences in envelope responses between PRE1 and PRE2 are not primarily driven by suppressive repetition effects but perhaps due to a consequence of change in task or cognitive processing demands. These distinctions together suggest that the effects of repetitions may be minimal in the current findings.

**Figure 4.**
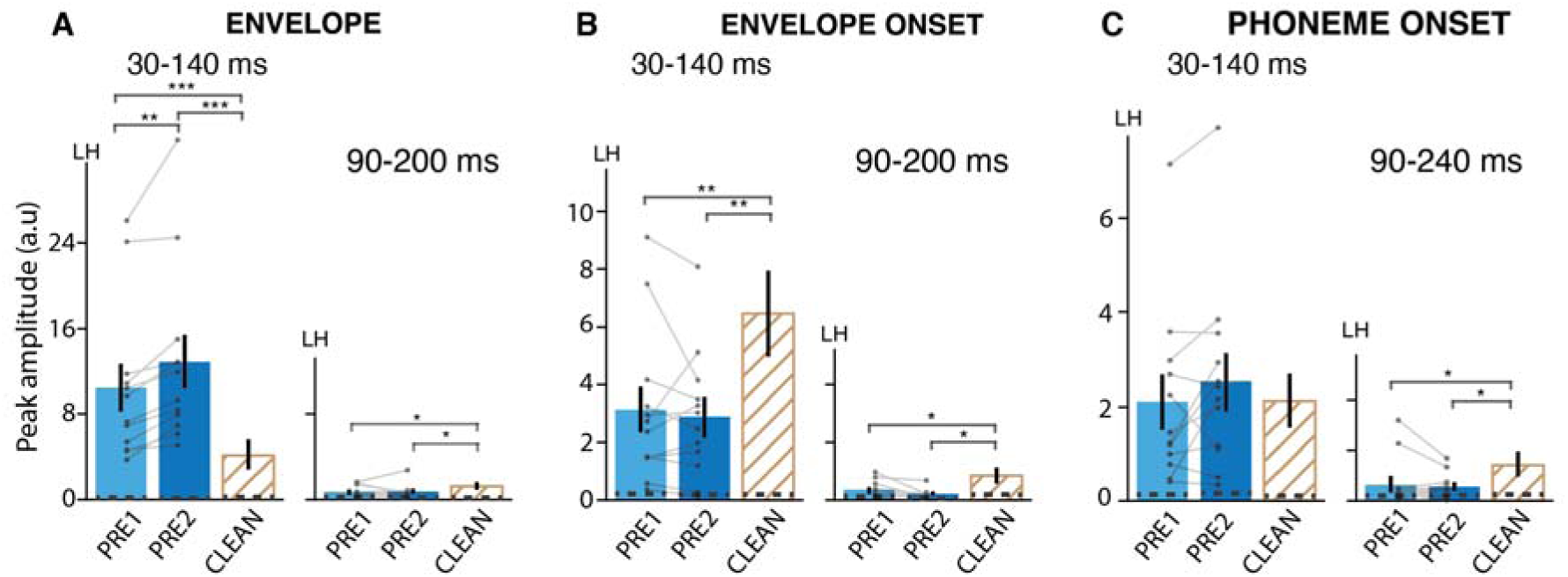
Neural responses to (A) Envelope, (B) Envelope onset, and (C) Phoneme onset in the control study. Other details as in Figure 2. Only the left hemisphere is shown (see supplemental Figure S2 for right hemisphere). The envelope onset early peak is stronger for PRE2 compared to PRE1. Differences between vocoded and clean are similar to those found in the main study.

### Additional analysis

Previous studies have indicated different roles for delta and theta band responses with respect to intelligibility and perceived clarity, with theta band responses showing links to clarity, and delta band responses to comprehension (45, 46). Therefore, we investigated whether the observed lexical-level changes with intelligibility in the low-frequency neural response (1-10 Hz) are specific to a neural frequency band. For this post-hoc analysis, word onset predicted neural response spectral power in each band was compared between speech conditions. Our results showed that both delta and theta band predicted response power increases from PRE to POST, with similar effect sizes (delta: *t_24_*= 2.50, *p* = 0.02, d = 0.51, theta: *t_24_* = 2.40, *p* = 0.02, d = 0.49). Thus, neither single interpretation of increased clarity vs. increased intelligibility can be given more prominence than the other.

In the present work, speech intelligibility is manipulated using a priming paradigm, where a perceptual pop-out effect modifies the perceived clarity of speech in addition to its intelligibility. It should be acknowledged that this approach is still subject to confounding factors, specifically those associated with predictive processing mechanisms and acoustical learning. Consistent with predictive coding theories, previous studies have seen that neural responses to degraded speech tend to be suppressed with better speech clarity (14, 23, 47). Conversely, it has also been proposed that activation of higher-order brain areas, which may exhibit less neural activity when speech clarity is compromised, can lead to a generally enhanced neural activation for intelligible speech (13, 23). Similarly, predictability in speech may amplify the synchronization of neural responses with speech, resulting in larger entrainment responses(46). Our findings align more closely with the latter perspective. Additionally, exposure to clear speech may induce changes in bottom-up processes that would facilitate the comprehension of degraded speech (13), perhaps including modifications to centrally maintained auditory filters at the subcortical and cortical level. Consequently, it is challenging to disentangle the precise neural mechanisms underpinning speech intelligibility from those intertwined with predictive processes.

Even though our analysis revealed effects of intelligibility on the word onset responses, no significant trend was observed between those neural measures and behaviorally-measured perceived speech clarity ratings within subjects. This might have been expected since the neural measures must be calculated across multiple trials, and minimizing the effects of individual trials because of the need to employ cross-validation, whereas the speech clarity did vary substantially from trial to trial (Supplemental Figure S3). It is unfortunate that estimating reliable TRFs from a single trial (in particular, estimating peaks with reliable amplitude and latency) is not feasible for only 20 seconds of data, either for acoustic or linguistic stimulus features. Similarly, the same limitation affects any analysis of the relationship between learning over trials and the word onset processing (i.e., early trails vs late trials).

In conclusion, we investigated the extent to which neural measures of lexical processing correspond to speech intelligibility while keeping the acoustical structure fixed. The neural measures associated with word onset processing, especially those time-locked at N400_TRF_ latencies (with N400 polarity), increased substantially between first exposure and after intelligibility-increasing priming. In contrast, auditory and phoneme onset responses are influenced only by the acoustics of the sensory input, not by intelligibility-boosting priming. It is crucial to exercise caution when interpreting auditory neural responses in contexts where the acoustics of the sensory input differ. Our key finding suggests that lexical segmentation responses increase in the same context as speech intelligibility and show engagement of top-down mechanisms. Together, these suggest that time locked neural responses associated with lexical segmentation may serve as an objective measure of speech intelligibility.

## Materials and Methods

### Participants

A total of 25 native English speakers (age range 18-32 yr, 15 males, 5 left-handed) participated in the main study. Data from two participants were excluded from the analysis due to excessive artifacts in the neural data. 12 native English speakers (age range19-27 yr, 5 males, 2 left-handed) participated in the control study. All participants reported normal hearing and no history of neurological impairments. The experimental procedures were approved by the University of Maryland institutional review board and all participants provided written informed consent before the experiment and were compensated for their time.

### Stimuli and Procedure

The stimuli were ∼20 s long excerpts (18 s – 26 s), sampled at 44.1 kHz, from the audiobook “The Botany of Desire” by Michael Pollan, narrated by a male speaker. Talker pauses greater than 400 ms were shortened to 400 ms and the excerpts were then low pass filtered below 4 kHz using a third order elliptic filter.

Noise-vocoded speech segments for each excerpt were generated using a custom python script. First, the frequency range 70-4000 Hz was divided into logarithmically spaced three bands (70 Hz - 432 Hz, 432 Hz - 1402 Hz, and 1402 Hz - 4000 Hz). For each band, an envelope modulated noise band was generated from band-limited white noise modulated by the envelope of the band passed speech signal. The envelope of the band passed speech signal was extracted using the half-wave rectification of the band passed signal followed by low pass filtering with a cutoff of 30 Hz. Finally, the modulated noise bands were summed, and the normalized volume was adjusted to match that of the original speech stimulus. Figure 1(A) shows example spectrogram plots for the original speech stimulus and vocoded speech stimulus, with the corresponding envelope amplitudes overlaid.

Subjects listened to a total of 36 trials that preserved the storyline. A trial consisted of noise-vocoded speech (PRE), followed by the same speech but in the original clear form (CLEAN) and then the second presentation of noise-vocoded speech (POST). All stimuli were presented diotically. At the end of each noise-vocoded passage, participants were asked to rate the perceived speech clarity (“How much could you follow the passage on a scale of 0 – 5?”; 0 - no words, 1 – a few words, 2 - definitely some words, 3 - lots of words but not most, 4 - more than half of all words, 5 - almost all words). The subjects were specifically instructed not to consider the CLEAN speech passage while making their decisions. This rating was used as a subjective measure of speech clarity. Intermittently, they were also asked to repeat back some of the words they could follow from the vocoded speech passage to ensure that the words they understood were actually from the passage. As a means of motivating the subjects to engage in the task, a comprehension-based question was asked at the end of the POST vocoded speech passage.

In the control study, stimuli, procedures and analyses were identical to those used in the main study, with the exception that the order of each CLEAN and POST vocoded speech were swapped. This resulted in a presentation order of PRE, POST and CLEAN for the control study, which are henceforth described as PRE1, PRE2, and CLEAN, respectively, to emphasize that the clean speech was not presented until after the second presentation of the vocoded speech.

### MEG data acquisition and preprocessing

Non-invasive neuromagnetic responses were recorded using a 160 channel whole head MEG system (KIT, Kanazawa, Japan), of which 157 channels are axial gradiometers and 3 magnetometers are employed as environment reference channels, inside a dimly lit, magnetically shielded room (Vacuumschmelze GmbH & Co. KG, Hanau, Germany) at the Maryland Neuroimaging Center. The data were sampled at 1 kHz along with an online low-pass filter with cut off frequency at 200 Hz and a 60 Hz notch filter.

During the task, subjects lay in supine position and were asked to minimize body movements as they listened and to keep their eyes open and fixate on a cross at the center of screen. Sound level was calibrated to ∼70 dB sound pressure level (SPL) using 500 Hz tones and equalized to be approximately flat from 40 Hz to 4 kHz. The stimuli were delivered using Presentation software (http://www.neurobs.com), E-A-RTONE 3 A tubes (impedance 50 Ω) which strongly attenuate frequencies above 4 kHz and E-A-RLINK (Etymotic Research, Elk Grove Village, United States) disposable earbuds inserted into the ear canals.

All data analyses were performed in mne-python 0.23.0 (48, 49) and Eelbrain 0.36 (50). Flat channels were excluded and the data were denoised using temporal signal space separation (TSSS) (51). Then the MEG data were filtered between 1 and 60 Hz using a zero-phase FIR filter (mne-python 0.23.0 default settings). Independent component analysis (ICA) (52) was then applied to manually remove artifacts such as eye movements, heart beats, muscle artifacts and singular artifacts. The cleaned data were low pass filtered between 1 and 10 Hz and downsampled to 100 Hz for further analysis.

### Neural source localization

The head shape of each participant was digitized using Polhemus 3SPACE FASTRAK three-dimensional digitizer. The position of the participant’s head relative to the sensors was determined before and after the experiment using five head-position indicator coils attached to the scalp surface and the two measurements were averaged. The digitized head shape and the marker coils locations were used to co-register the template FreeSurfer “fsaverage” (53) brain to each participant’s head shape using rotation, translation and uniform scaling.

A source space was formed by four-fold icosahedral subdivision of the white matter surface of the fsaverage brain, with all source dipoles oriented perpendicularly to the cortical surface. The source space data and the noise covariance estimated from empty room data were used to compute the inverse operator using minimum norm current estimation (54, 55). The analysis were restricted to frontal, temporal and parietal brain regions based on the ‘aparc’ FreeSurfer parcellation (56). Excluded brain regions are shaded in dark gray in the brain plots (Figure 2, Figure 3).

### Predictor variables

The speech signal was transformed into unique feature spaces to represent different levels of the language hierarchy. These feature-based model predictors can be categorized into three main groups: 1. Acoustic (acoustic envelope and acoustic onsets), 2. sub-lexical (phoneme onset), 3. lexical (word onset, contextual word surprisal). All predictor variables were downsampled to 100 Hz.

### Acoustic Properties

The acoustic envelope predictor reflects instantaneous acoustic power, and the acoustic onset reflects the salient transients, of the speech signal. Both of these continuous representations were computed via a simple model of the human auditory system using gammatone filters with Gammatone Filterbank Toolkit 1.0 (57). A filterbank-based broad-band envelope extraction method was used (instead of the conventional broad-band envelope extraction method: absolute value of the Hilbert-transformed signal), as it has been shown that the filterbank-based broad-band envelope increases the neural tracking of the speech envelope (58). First, the gammatone spectrogram was generated with cut-off frequencies from 20 to 5000 Hz, 256 filter channels and 0.01 s window length. Each frequency band was then resampled to 1000 Hz and transformed to log scale. Then the envelope spectrogram was averaged across 256 channels, resulting in a broad-band temporal acoustic envelope predictor.

The acoustic onset representations were computed on the gammatone acoustic envelope spectrogram, by applying an auditory edge detection algorithm (59). Similar to the acoustic envelope, the onset spectrogram was also averaged across frequency bands.

In order to estimate differences in the acoustic feature predictors between clean and vocoded speech, we used the linear-correlation coefficient *r*. We observed a strong positive correlation between vocoded and clean passages for the envelope (*r* = 0.92, *p* < 0.001), as expected from the method used to construct the vocoded speech. However, this correlation was smaller for the envelope onset (*r* = 0.46, *p* < 0.001), since acoustic onsets are less well preserved than the explicitly copied envelope temporal modulations in the noise-vocoded speech (60).

### Sub-lexical Properties

The Montreal Forced Aligner (61) was used to align the speech acoustics with the words and phonological forms from a pronunciation dictionary. The CMU Pronouncing Dictionary (http://www.speech.cs.cmu.edu/cgi-bin/cmudict), excluding the stress information, was used as the pronunciation lexicon. The pronunciation lexicon, transcriptions and audio file were aligned using the ‘english’ pretrained acoustic model. The annotations for phoneme and word onsets were visualized in PRAAT (62) and manually adjusted appropriately. The phoneme onsets predictor was represented as the impulses at the onset of each phoneme.

### Lexical Properties

The word onsets were represented as unit impulses at the onset of each word.

Contextual word surprisal was estimated using an open source transformer-based (63) large language model (GPT-2) (42). We used gpt2-large pretrained models implemented in the Hugging Face environment (64). The transcripts for each passage were preprocessed by removing punctuation and capitalization (places and names were retained). The model first tokenizes the words, where the tokens could represent either words or sub-words, and then fed into the model. The model outputs activation at each of the 36 layers in the network and we used the final layer for the word surprisal calculation. The final layer includes prediction scores given the previous context for each word in the passage over the vocabulary token. Here the ‘context’ refers to all the preceding tokens or sequence of tokens at least 1024 tokens long. The prediction scores were SoftMax transformed to compute the probability. The current word probability was computed by the probability of the corresponding token and for the words spanning over multiple tokens, the word probability was computed by the joint probability of the tokens. Contextual word surprisal was computed as the - log_2_(P_WOTd_|context) and represented as an impulse at each word onset scaled by the corresponding model surprisal of that word.

Despite the differences in acoustical features between the vocoded and clean speech passages, sub-lexical and lexical feature time series were kept unchanged.

#### Computational model

A linear forward modeling approach using Temporal response functions (TRFs) was used to analyze the phase-locked neural responses to various speech features simultaneously (17, 65). Analogous to the conventional event related potentials (ERP), TRFs estimate how the brain responds to speech features over time, or from the signal processing viewpoint, the brain’s impulse response to any given speech feature. In contrast to ERPs, that rely on averaging (perhaps) hundreds of short responses to estimate the brain responses to a given stimulus, TRF analysis allows to determine the brain responses to long-duration continuous speech. Critically, TRFs can also model simultaneous responses to multiple speech features (1), multivariate form of the TRFs (mTRF).

Here, TRFs were estimated using the boosting algorithm, which minimizes the l1 error between the measured and predicted source current time course, over the time lags -200 ms to 800 ms (using a basis of 50 ms width Hamming windows). Fourfold cross-validation (two training sets, one testing set and one validation set) was used to prevent overfitting and improve the generalized performance. Prediction accuracy was estimated as the explained variance of the TRF model. The subsequent statistical analysis was performed using each subject’s average TRFs and prediction accuracies across all cross-validation folds. In sum, optimal TRFs were estimated for each subject, speech condition, and each source current dipole including multiple speech features simultaneously.

Prior estimating TRFs, predictors (speech features) and neural responses were z-score normalized, so that they can be compared between subjects and conditions. To visualize the TRFs over ROI (temporal, frontal and parietal) as a single time series with the current direction, TRFs over ROIs were simplified using principal component analysis (PCA). Examples of the first principal component are shown in Figure 2(A) and Figure 3(A).

#### TRF peak amplitude extraction

The TRFs showed prominent peaks at different latencies. Based on the TRFs across subjects, we identified time windows of interest and polarity for each peak; envelope TRF: P1 (30-140 ms) and N1 (90-200 ms), envelope onset TRF: P1 (30-140 ms) and N1 (90-200 ms), phoneme onset TRF: P1 (30-140 ms) and N1 (90-240 ms), word onset and contextual word surprisal TRFs: P1 (50-300 ms) and N1 (330-650 ms), where P1 and N1 represent the polarity of the current estimate, positive and negative respectively, in the time windows specified above. The average TRF response for each subject and condition were obtained as the sum of absolute current dipoles across the ROI. TRF peaks for each subject and condition were picked by searching for the maximum peak in the average TRF, that aligned with the current direction from the original source TRFs. The polarity of the source TRFs were determined by the current direction relative to the cortical surface at the transverse temporal region. If none of the peaks satisfied the polarity constraint, the minimum of the average TRFs in the given time window was used as the peak amplitude.

To aid the TRF peak evaluations, TRF peak amplitude noise floor was measured on noise model TRFs. The noise model TRFs were generated by mismatching the predictors and neural data and estimating the TRFs with the same parameters used in the TRF estimation. These noise model TRFs were then subjected to the same peak picking algorithm as described above to measure the noise model TRF peak amplitudes for each subject. The TRF peak amplitude noise floor for each peak and condition is calculated as the mean noisy peak amplitude across subjects, and represented by dashed lines in the bar plots (See Figure 2 and Figure 3). This noise floor serves as a reference for evaluating the significance of the observed TRF peaks.

#### Quantification and Statistical Analysis

Statistical analysis was performed in R version 4.0 (66) and Eelbrain. The significance level was set at a = 0.05.

Linear mixed effect model (LMEM) analysis was performed to evaluate the trends in behavioral measures. For the LMEM analysis, the *lme4* (version 1.1-30) (67), *lmerTest* (version 3.1-30) (68) and *buildmer* (version 2.4) (69) packages in R were used. The best fit model from the full models were determined using the *buildmer* function. The assumptions of mixed effect modelling, linearity, homogeneity of variance, and normality of residuals, were checked per each best fit model based on the residual plots. Reported effect sizes represent the changes in the dependent measure when comparing one level of an independent variable to its reference level. *p*-values were calculated using Satterthwaite approximation for degrees of freedom (70, 71).

To compare the predictive power of two models, such as the full model vs reduced or PRE vs POST, the difference in explained variability at each source dipole was calculated. Significant differences between the two models were tested while controlling for multiple comparisons, using one-tailed paired-sample *t*-test with threshold-free cluster enhancement (TFCE) (72) and with a null distribution based on 10000 random permutations of the condition labels. The largest *t*-value and the corresponding *p* value are reported in results. To visualize the model fits in meaningful units, they were scaled by the largest explanatory power of the full model across subjects, allowing expression as a % of the full model. The model comparison plots (predictive power of each feature) reported in Figure 2 and Figure 3 are masked by significance to emphasize how each feature contributes to the model.

Lateralization tests were performed to estimate any hemispheric asymmetry in speech feature processing. To accomplish this, the predictive power was first transferred to a common space by morphing the source data to the symmetric “fsaverage_sym” brain, followed by morphing the right hemisphere to the left hemisphere. Once the data were put in the common space, a two-tailed paired sample *t*-test with TFCE was used to test for significant differences between the left and right hemispheres.

TRF peak amplitudes between conditions were compared using paired-sample *t*-test. TRF peak amplitude comparisons between main study and control study were performed using two-sample randomization (permutation) test using *EnvStats* (Version 2.7) (73) R package. The effect sizes for paired or independent sample *t*-tests were calculated using Cohen’s d (74) (d), where d = 0.2 indicates a small effect, d = 0.5 indicates a moderate effect, and d = 0.8 indicates a large effect.

To test the changes in spectral power in delta and theta bands, the predicted responses were subjected to power spectral analysis. Spectral power estimates were averaged across trials, frequency (delta: 1-4 Hz and theta: 4-8 Hz), and source data. Spectral power estimates were compared across conditions using a paired-sample *t*-test.

The number of subjects for the control study was determined using a power analysis (*power.t.test* function in R). This analysis was focused on the word onset late responses to detect an effect of priming. Analysis indicated that a sample size of nine subjects is sufficient to detect the desired effect with a power of 0.8. We instead included twelve subjects in the control experiment, exceeding the minimum necessary sample size determined by the power analysis.

## Handedness

In our study, data from all participants were analyzed, regardless of their handedness, as we did not have any specific hypothesis related to handedness. Nevertheless, to examine any potential influence, we compared the results before and after excluding the data from the five left-handed participants. This comparison revealed that exclusion of left-handed participants did not affect any of the findings of significance for PRE vs POST or PRE vs CLEAN comparisons. For the POST vs CLEAN comparison, no findings of significance were affected except for the late word onset responses, where significance was lost when left-handed subjects were excluded.

## Data availability

The MEG data, behavioral responses and codes to reproduce the results in this paper will be shared once the paper is accepted

## Author Contributions

I.M.D.K., J.P.K, and J.Z.S. conceived and designed the research; I.M.D.K. collected data; I.M.D.K. and J.Z.S. analyzed data; I.M.D.K. drafted manuscript; I.M.D.K., J.P.K and J.Z.S. edited and revised manuscript.

## Acknowledgments

This work was supported by the National Institutes of Health grant R01-DC019394 and National Science Foundation grant SMA 1734892. We thank Philip Resnik and Shohini Bhattasali for their assistance in choosing context-based linguistic feature and Ciaran Stone for excellent technical assistance.

## Supplementary Information

**Table S1.**
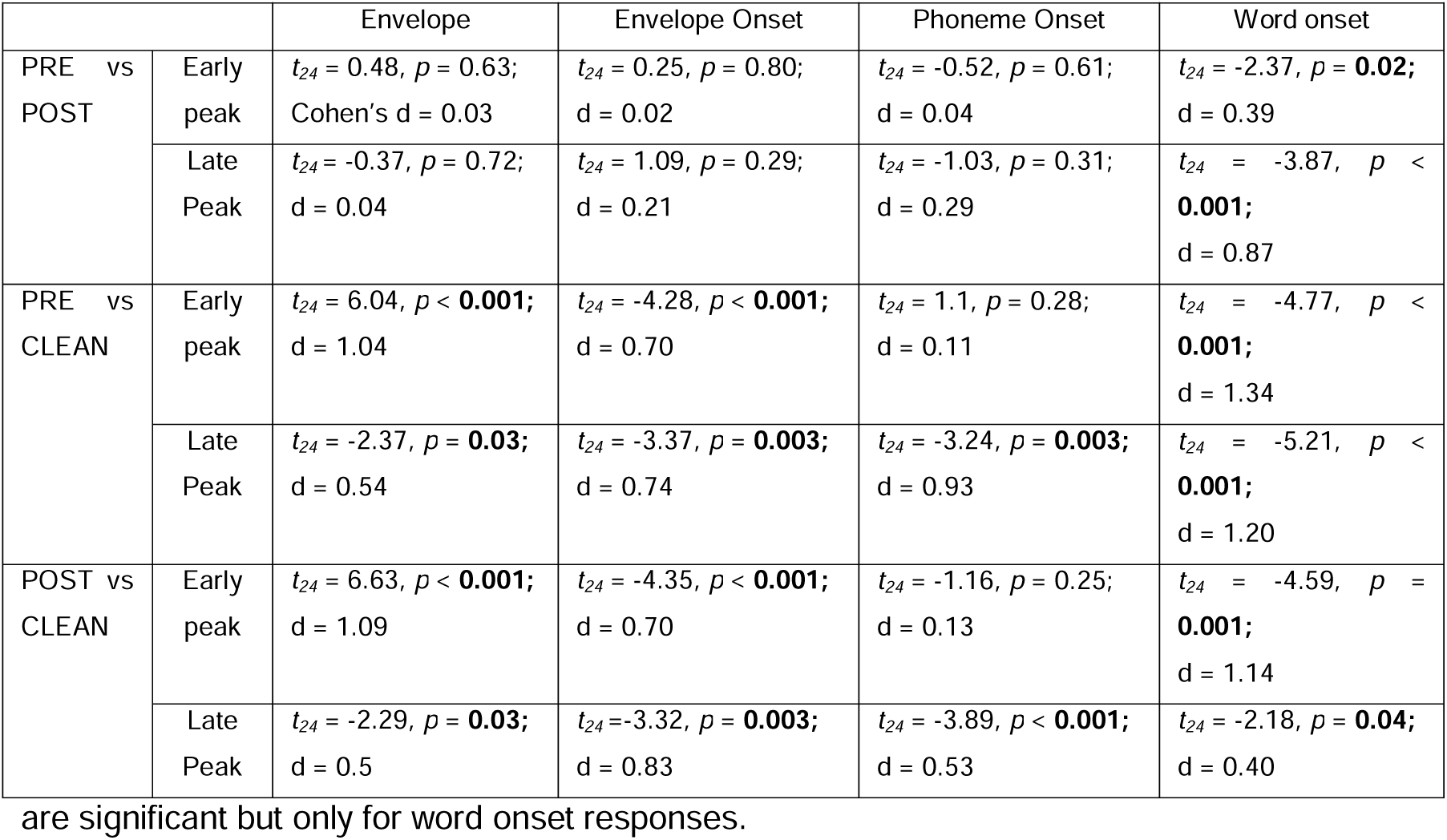
Summary Statistics for TRF peak amplitude comparisons between PRE vs POST, PRE vs CLEAN, and POST vs CLEAN and for both early and late peak (paired t values, p values, and Cohen’s d for effect size) in the main study. Differences between CLEAN and either PRE or POST are significant except for phoneme onset early peak. Differences between PRE and POST are significant but only for word onset responses.

**Table S2.** Summary Statistics for TRF peak amplitude comparisons between PRE1 vs PRE2, PRE1 vs CLEAN, and PRE2 vs CLEAN and for both early and late peak (paired t values, p values, and Cohen’s d for effect size) in the control study. Differences between CLEAN and either PRE1 or PRE2 are significant for early peaks except for phoneme onset. Almost all differences between PRE1 and PRE2 are not significant.

|  |  | Envelope | Envelope Onset | Phoneme Onset | Word onset |
| --- | --- | --- | --- | --- | --- |
| PRE1 vs PRE2 | Early peak | $t_{11} = -4.23, p = \mathbf{0.001}$; Cohen's d = 0.31 | $t_{11} = 0.55, p = 0.59$; d = 0.09 | $t_{11} = -1.76, p = 0.11$; d = 0.23 | $t_{11} = 1.01, p = 0.34$; d = 0.22 |
| | Late Peak | $t_{11} = 0.08, p = 0.94$; d = 0.02 | $t_{11} = 1.79, p = 0.10$; d = 0.49 | $t_{11} = 0.66, p = 0.52$; d = 0.14 | $t_{11} = 0.35, p = 0.74$; d = 0.06 |
| PRE1 vs CLEAN | Early peak | $t_{11} = 4.41, p = \mathbf{0.001}$; d = 1.01 | $t_{11} = -3.90, p = \mathbf{0.002}$; d = 0.87 | $t_{11} = -0.19, p = 0.85$; d = 0.02 | $t_{11} = -4.98, p < \mathbf{0.001}$; d = 1.63 |
| | Late Peak | $t_{11} = -2.32, p = 0.04$; d = 0.83 | $t_{11} = -2.31, p = 0.04$; d = 0.84 | $t_{11} = -2.84, p = 0.02$; d = 0.63 | $t_{11} = -5.11, p = \mathbf{0.009}$; d = 1.76 |
| PRE2 vs CLEAN | Early peak | $t_{11} = 4.77, p < \mathbf{0.001}$; d = 1.27 | $t_{11} = -3.43, p = \mathbf{0.006}$; d = 0.96 | $t_{11} = 1.39, p = 0.19$; d = 0.21 | $t_{81} = -4.71, p < \mathbf{0.001}$; d = 1.84 |
| | Late Peak | $t_{11} = -2.12, p = 0.06$; d = 0.78 | $t_{11} = -2.88, p = 0.02$; d = 1.07 | $t_{11} = -2.50, p = 0.03$; d = 0.80 | $t_{11} = -5.26, p < \mathbf{0.001}$; d = 1.86 |

**Figure S1.**
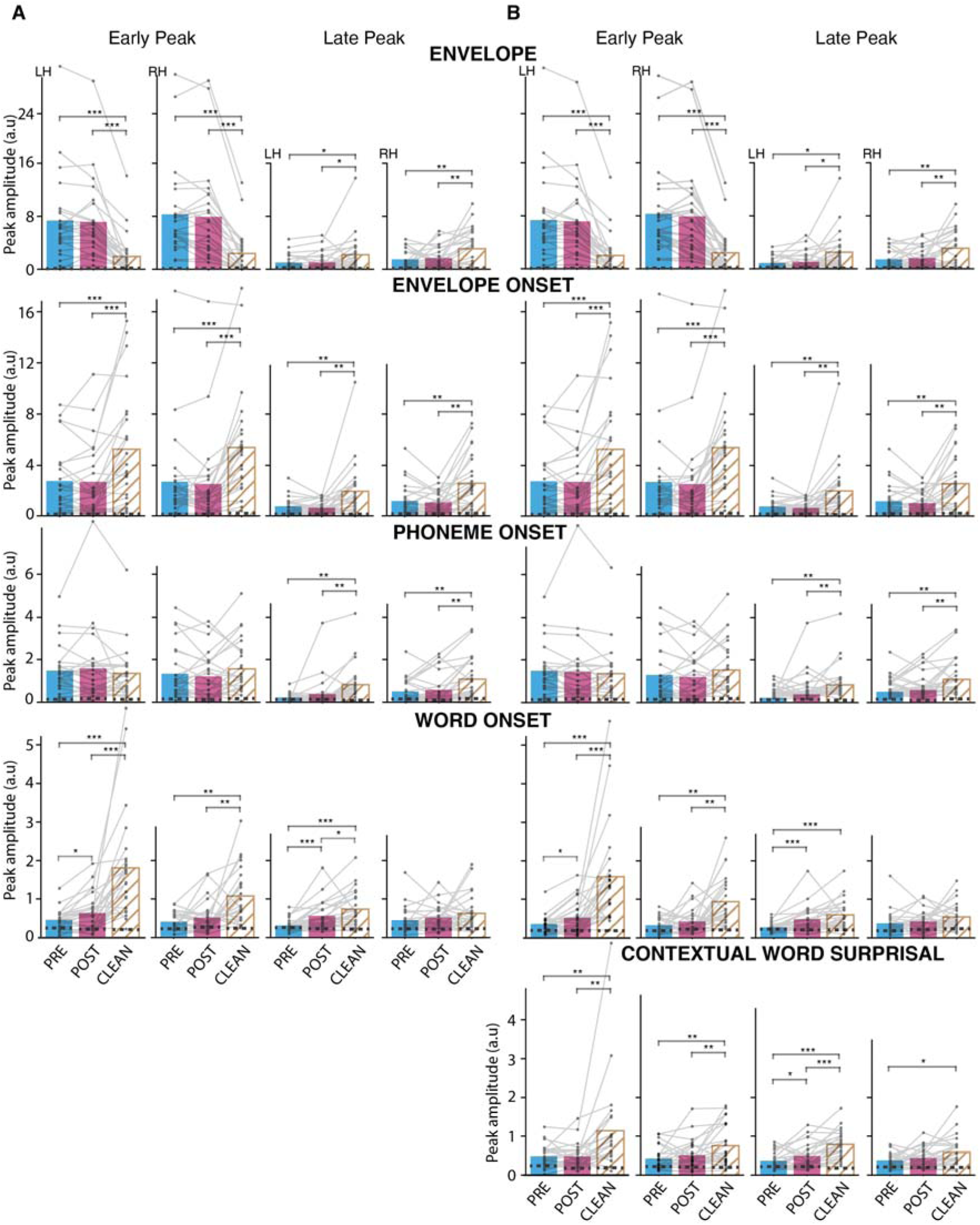
TRF peak amplitude comparison by speech condition for envelope, envelope onset, phoneme onset, word onset and contextual word surprisal, for early and late peaks, and by hemisphere (A) for the same TRF model as in Figure 2 and (B) for another model including contextual word surprisal. The dashed lines within the bars represent the noise floor. Other details as in Figure 2. Similar to the main model, almost all differences between CLEAN and either PRE or POST are significant, except for phoneme onset early peak. Differences between PRE and POST are significant only for word onset responses and contextual word surprisal late responses in the left hemisphere. Adding contextual word surprisal into the model does not substantially change significant intelligibility effects observed in word onset responses. *p<0.05, **p<0.01, ***p<0.001

**Figure S2.**
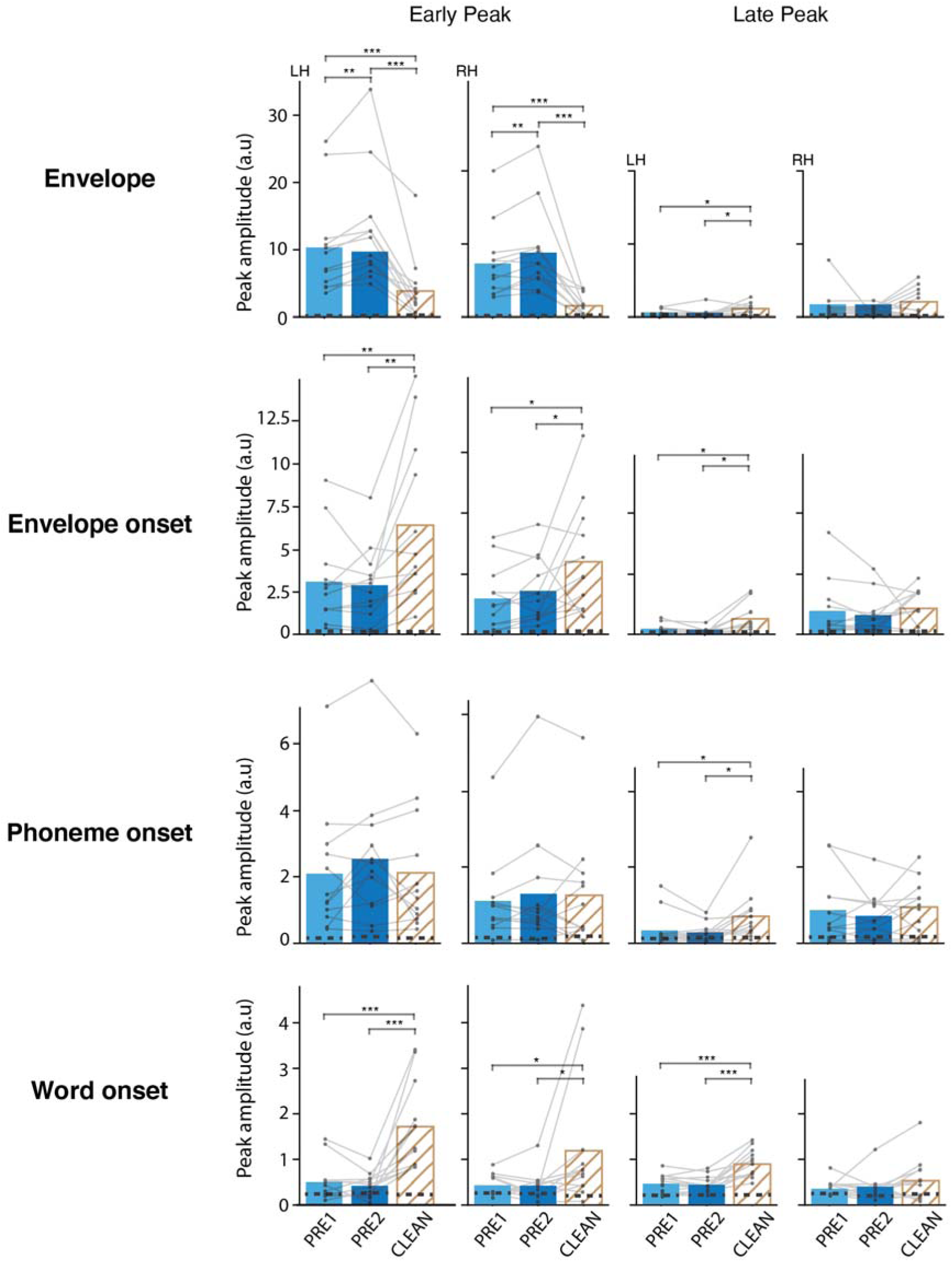
Control study TRF peak amplitude comparison by speech condition for envelope, envelope onset, phoneme onset, and word onset, for early and late peaks, and by hemisphere. Other details as in Figure 2. Differences between PRE1 and PRE2 are significant only for early envelope responses in both hemispheres. Differences between vocoded and CLEAN are similar to those observed in the main study.

**Figure S3.**
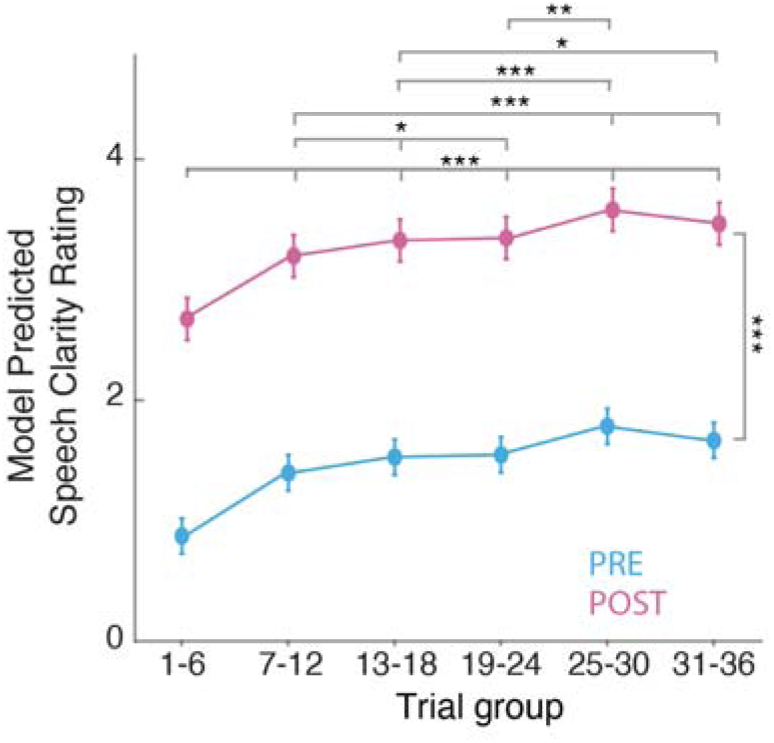
Speech clarity rating change over trials. Trials were combined using a tumbling window of 6 trials. Linear Mixed Effects Model (clarity rating ∼ 1 + condition + trial group (1+condition|Subject)) predicted clarity rating ± SE change over trial group. As no significant interaction was found between condition and trial group, the model’s trends over trials for both PRE and POST are parallel. Overall, speech clarity improves over trials for both PRE and POST vocoded speech (trials 1-6 as the reference, trials 7-12 = 0.53, SE = 0.06, p < 0.001, trials 13-18 = 0.66, SE = 0.06, p < 0.001, trials 19-24 = 0.68, SE = 0.06, p < 0.001, trials 25-30 = 0.91, SE = 0.06, p < 0.001, trials 31-36 = 0.79, SE = 0.06, p < 0.001) and then level off (trials 19-24 to trials 25-30 = 0.15, SE = 0.06, p = 0.33). *p<0.05, **p<0.01, ***p<0.001

